# Cellular endosomal potassium ion flux regulates arenavirus uncoating during virus entry

**DOI:** 10.1101/2023.06.23.546275

**Authors:** Amelia B. Shaw, Hiu Nam Tse, Owen Byford, Grace Plahe, Alex Moon-Walker, Erica Ollmann-Saphire, Sean P. J. Whelan, Jamel Mankouri, Juan Fontana, John N. Barr

## Abstract

Lymphocytic choriomeningitis virus (LCMV) is a model arenavirus that causes fatalities within immunocompromised populations. To enter cells, the LCMV envelope fuses with endosomal membranes, for which two requirements are low pH and interaction between LCMV GP spike and receptor CD164. LCMV subsequently uncoats, where genome-associated NP separates from Z matrix. To further examine LCMV entry, an siRNA screen identified K^+^ channels as important for LCMV infection, and pharmacological inhibition confirmed K^+^ involvement during entry. We tracked incoming virions along their entry pathway under physiological conditions, where uncoating was signified by separation of NP and Z. In contrast, K^+^ channel blockade, prevented uncoating, trapping virions within Rab7 and CD164-positive endosomes, identifying K^+^ as a third LCMV entry requirement. K^+^ did not increase GP/CD164 binding, thus we suggest K^+^ mediates uncoating by modulating NP/Z interactions within the virion interior. These results suggest repurposing licensed K^+^ channel inhibitors represents a potential anti-arenaviral strategy.

## INTRODUCTION

The *Arenaviridae* family within the *Bunyavirales* order of enveloped segmented RNA viruses currently contains around 50 species grouped into four genera, *Antennavirus*, *Hartmanivirus*, *Mammarenavirus* and *Reptarenavirus*^1, 2^. Mammarenaviruses can be segregated into New World (NW) and Old World (OW) groups based on phylogeny, with both groups comprising several human-infecting arenaviruses responsible for fatal haemorrhagic fevers. Mammarenaviruses are associated with rodent hosts that act as stable reservoirs. Human disease typically results from spill-over events, in areas where human and rodent habitats coincide, but also occurs in nosocomial settings, providing a high risk to healthcare workers^3, 4^. Important human pathogens from the NW group include Junín (JUNV), Machupo (MACV) and Guanarito viruses, alongside Lassa and Lujo viruses (LASV and LUJV, respectively), from the OW group. The OW lymphocytic choriomeningitis virus (LCMV) is associated with neurologic disease in immunocompromised individuals and is a well-known cause of miscarriages but is largely non-pathogenic to immunocompetent humans^5, 6^. As such, LCMV represents a tractable model system for studying arenavirus molecular and cellular biology and acts as the prototype of the entire *Arenaviridae* family. Options for mitigation of arenavirus disease are limited, with very few effective preventative or therapeutic measures currently available.

The mammarenavirus genome comprises small (S) and large (L) RNA segments that use an ambi-sense strategy to encode a total of four proteins; the S segment encodes the nucleocapsid protein (NP) and the glycoprotein precursor (GPC), whereas the L segment encodes both the large protein (L), which is the catalytic component of the viral RNA-dependent RNA-polymerase, and the Z matrix protein (Z). GPC is co- and post-translationally cleaved by cellular proteases, yielding the stable signal peptide (SSP), GP1 and GP2, which exist together on the viral surface as tripartite trimers^7, 8^. Arenavirus entry into cells is directed by GP1, responsible for receptor binding, and GP2, which promotes fusion between host and viral membranes^9^. GP1 binds various cell surface receptors such as α-dystroglycan (LCMV and LASV)^10^, neuropilin-2 (LUJV)^11^, and transferrin receptor (JUNV and MACV)^12–14^ although other alternate receptors have been implicated including Tyro3/Axl/Mer and members of the T-cell immunoglobulin (TIM) and mucin receptor families^15, 16^. Subsequently, arenaviruses are internalized through receptor-mediated endocytosis, to traffic through the endocytic system. The changing biochemical composition of maturing endosomes induces structural changes to the GPC to trigger at least two critical events; the first is a receptor switch mechanism, in which GP1 binds an alternate endosomal-resident receptor, with LASV interacting with LAMP-1^17, 18^, LUJV with CD63^11^ and LCMV requiring CD164^19, 20^. The second is a conformational rearrangement of GP2, which exposes a fusion loop that mediates viral and cellular membrane fusion to release the genomic segments into the cytosol.

Endosomal acidic pH is a well-characterised trigger of enveloped virus entry, activating viral spikes resulting in fusion between virion and endosomal membranes at characteristic locations within the endocytic system. However, it is emerging that other ions also act as entry triggers; for example, binding of calcium has been shown to promote entry of togaviruses^21^, coronaviruses^22–24^ and filoviruses^25^, whereas potassium ions (K^+^) are required for uncoating of influenza virus by promoting separation of its matrix and ribonucleoprotein (RNP) components^26^. Within the *Bunyavirales* order, we recently showed K^+^ expedited entry of Bunyamwera peribunyavirus (BUNV) and Hazara nairovirus (HAZV) by inducing structural changes in their virion spikes^27–29^, promoting fusion between the virus and its target membranes^28, 30^.

Here, we explored the ionic requirements for arenavirus multiplication. We performed an siRNA-mediated gene knock-down screen that showed arenavirus growth depended on several cellular K^+^ channels. Pharmacological inhibition corroborated these findings, with time-of addition studies showing K^+^ channel activity was required during virus entry. K^+^ channel blockade trapped virions within Rab7-positive late endosomal compartments where virion uncoating was prevented despite the presence of second receptor CD164. Using a previously established CD164/GP1 binding assay, we showed the role of K^+^ in virion uncoating was not to promote this receptor interaction. Instead, we propose K^+^ modulates interactions between virion components within the virion interior.

Taken together, these findings reveal an important role for K^+^ during LCMV entry and the opportunity for existing, licensed K^+^ ion channel inhibitors to be repurposed as a new, pharmacologically safe and broad-ranging therapeutic strategy for arenavirus-mediated disease.

## RESULTS

### Generation of an eGFP-expressing LCMV variant

To investigate the dependence of LCMV on endosomal ion flux, an initial aim of this study was to perform a comprehensive siRNA screen of host cell ion channels involved in LCMV multiplication. To facilitate this, we generated a recombinant LCMV (rLCMV) expressing eGFP (rLCMV-eGFP) to permit live-cell monitoring of virus-specific gene expression.

First, to rescue wild-type rLCMV (rLCMV-WT), cDNAs pUC57-LCMV-S and pUC57-LCMV-L were designed to express wild-type LCMV-Armstrong (clone 13 derivative) S and L segment sequences, respectively (Supplementary Fig. 1A). To confirm successful rescue, a silent *Xh*oI site was introduced into pUC57-LCMV-S, to permit distinction of rescued rLCMV from non-recombinant LCMV stocks (Supplementary Fig. 1B-E). To generate the rLCMV-eGFP variant, pUC57-LCMV-S was modified by insertion of a porcine teschovirus-1 2A peptide linker (P2A) sequence^31–33^ between eGFP and LCMV NP ORFs (Supplementary Fig. 2A). All viruses were rescued in BSR-T7 cells, with rescue confirmed by detection of NP by western blot analysis of both transfected and infected cell lysates (Supplementary Fig. 2B).

### rLCMV and rLCMV-eGFP exhibit similar growth kinetics

We next compared the multi-step growth kinetics of rLCMV-WT and rLCMV-eGFP to assess the fitness impact of the eGFP ORF and P2A linker. BHK-21 cells were infected with each virus at an MOI of 0.001 and released virus was measured at 24 hr intervals by immuno-staining focus forming assay using NP antisera. This analysis revealed multiplication of rLCMV-WT and rLCMV-eGFP were similar, with less than 1-log difference between titres across the time course (Fig. 1A). Comparison between immuno-stained and fluorescent foci showed eGFP expression was an accurate surrogate marker for the detection of rLCMV-infected cells (Fig. 1B), with benefits of not requiring antibody staining and allowing tracking infection in live cells.

**FIGURE 1.**
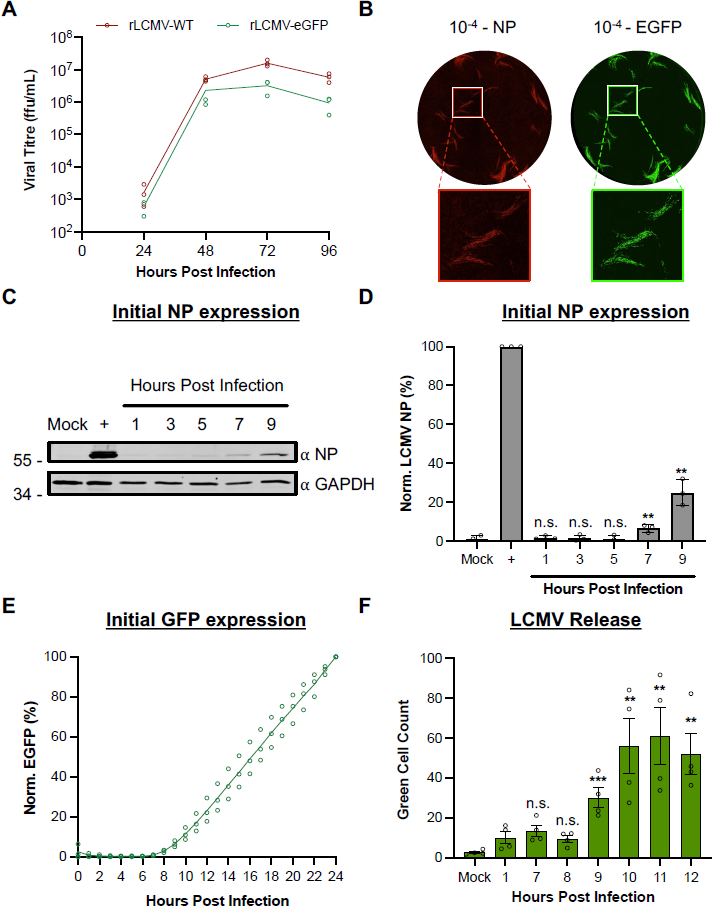
Comparison of replication kinetics of wildtype LCMV and eGFP-expressing LCMV. (A) BHK-21 cells were infected in triplicate (n=3) with either rLCMV-WT or rLCMV-eGFP at a MOI of 0.001. Supernatant samples were collected every 24 hours and were subsequently titred by focus forming assay. The average titre of each timepoint was plotted with standard error. (B) Focus forming assays for LCMV were performed by infecting BHK-21 cells with a serial dilution of the virus supernatant and incubating for 3 days. For rLCMV-WT, the focus forming assay was fixed, permeabilised and stained for LCMV NP by indirect immunostaining and imaged using the Incucyte S3. For rLCMV-eGFP, the focus forming assays were imaged without fixing. (C-D) To identify initial synthesis of NP, SH-SY5Y cells were infected with rLCMV-WT and lysates were collected at subsequent hours post infection for western blot (C) and densitometry analysis (D). This was plotted as a percentage of LCMV NP expression at 24 hours post infection, with standard error (n=3). Results were analysed by Student’s t-test whereby n.s. = p>0.05, * = p<0.05; ** = p<0.01; *** = p<0.001, comparing the timepoints to NP expression at 24 hours. (E) To examine timepoints of eGFP expression, SH-SY5Y cells were infected with rLCMV-eGFP and total integrated intensity of eGFP expression was measured every hour using the incucyte S3 and plotted as a percentage of eGFP expression at 24 hours, with standard error (n=3). (F) SH-SY5Y cells were infected with rLCMV-eGFP at an MOI of 0.2 for one hour at 37 ℃, after which the virus supernatant was removed, and the cells extensively washed with PBS. Fresh media was added to the cells and incubated until the specified timepoint to transfer the supernatant to fresh cells, which were then imaged, using the Incucyte S3, 24 hours after the time of transfer. The average number of eGFP-expressing cells counted (Green Cell Count), resulting from four experimental repeats, is plotted with standard error. Results were analysed by Student’s t-test whereby n.s. = p>0.05, * = p<0.05; ** = p<0.01; *** = p<0.001, comparing timepoints to 1 hour post infection.

To better characterise NP and eGFP expression kinetics, human neuronal SH-SY5Y cells, relevant as neuronal cells are targeted during LCMV host infection, were infected with either rLCMV-WT or rLCMV-eGFP at an MOI of 0.1 and expression was measured at multiple hours post infection (hpi). For rLCMV-WT, NP expression was first visible at 7 hpi (Fig. 1C-D). For rLCMV-eGFP, new eGFP expression was quantified through measuring the total integrated intensity of the eGFP (TIIE) signal, first detected at 8 hpi (Fig. 1E), suggesting the gene expression kinetics of these viruses were similar. Finally, to determine the time taken for a single replication cycle of rLCMV-eGFP, we performed a virus release assay. Cells were infected with rLCMV-eGFP, and at specified time points post-infection, all supernatant was transferred to fresh cells. The presence of infectious virus was then detected by incubating these fresh cells for 24 h, at which time the green cell count was taken to determine the number of eGFP-expressing cells. This analysis showed that virus was first released from cells at 9 hpi (Fig. 1F), indicating this was the minimum time required for a complete rLCMV-eGFP multiplication cycle in these cells.

### Cellular K^+^ channels are required for LCMV replication

Next, we used rLCMV-eGFP to identify host cell ion channels that play a role during LCMV growth. To achieve this, we used a curated siRNA library comprising three unique RNA sequences targeting over 150 ion channel genes and assessed the ability of rLCMV-eGFP to multiply in the context of gene knockdown. Following reverse transfection of each unique siRNA, SH-SY5Y cells were infected with rLCMV-eGFP at an MOI of 0.1 for 16 h, after which the TIIE signal was quantified as a marker of successful rLCMV-eGFP infection (Fig. 2A). As previously determined, initial eGFP expression is first detected at 8 hours, with rLCMV-eGFP first released from cells at 9 hpi (Fig. 1E-F). While virus will have been released from initially infected cells within the 16 hpi timepoint, eGFP expression in any newly infected cells will not be detectable. Thus, quantification of the eGFP signal recorded at 16 hpi would allow the maximum expression of eGFP, while only reflecting the influence of siRNA knockdown on rLCMV-specific activities up to and including eGFP expression within the first round of infected cells. The influence of siRNA knockdown on later events in the growth cycle, including virus assembly, budding and infection of new cells would not be represented in the TIIE signal.

**FIGURE 2.**
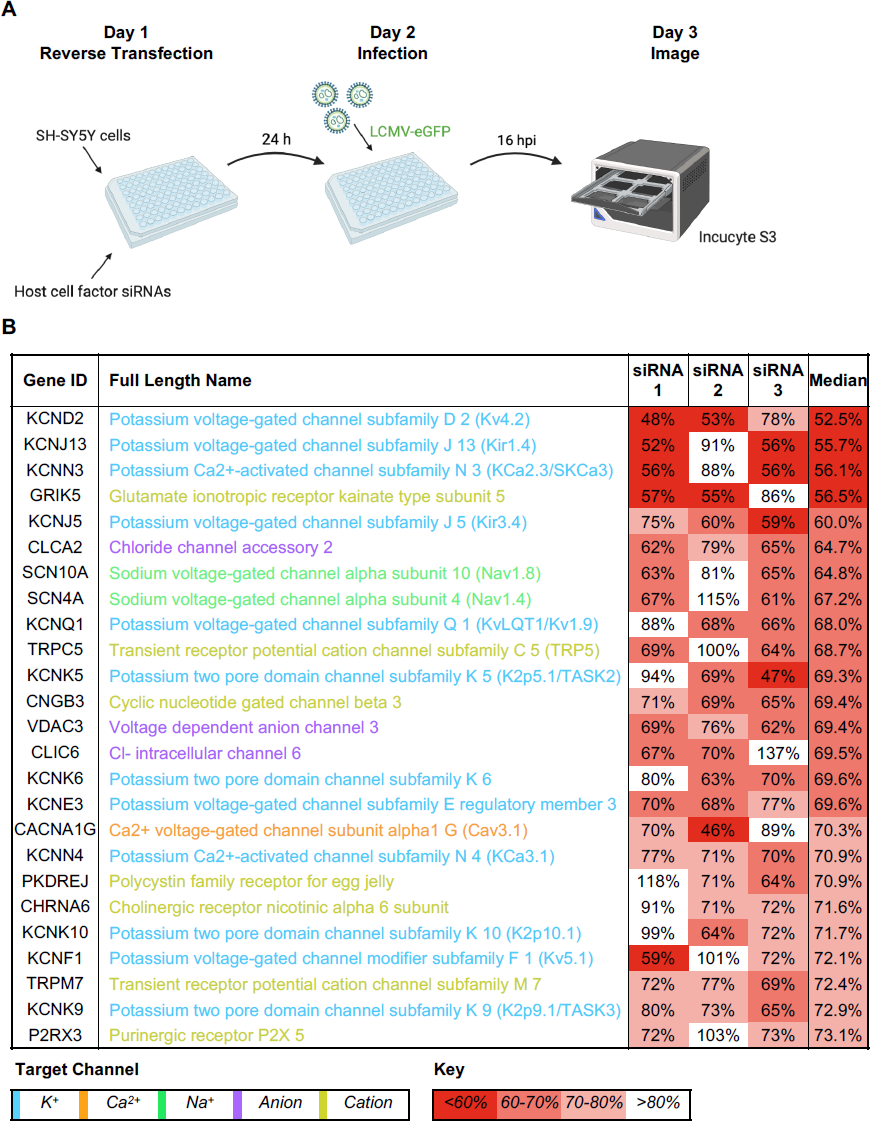
Assessment of the requirement of ion channels in the LCMV lifecycle using a siRNA screen. (A) Schematic depiction of the siRNA screening protocol. SH-SY5Y cells were reverse-transfected with siRNAs that targeted 176 cellular ion channel genes and then infected with rLCMV-eGFP. At 16 hours post infection (hpi), the cells were scanned using the incucyte S3 for expression of eGFP. (B) The top 25 gene targets which resulted in the highest knockdown of eGFP expression at 16 hpi have been shown. Percentage knockdown has been shown as the median of the average knockdown percentage of each of three individual siRNA targeting the particular gene. Each percentage represents the mean of four experimental repeats. The channel families have been classified into colour as shown by the target channel box. The percentage knockdown has been colour coded by shades of red to indicate the level of knockdown, with more red representing over 60 % knockdown of expression and white representing less than 80 % knockdown of expression.

The effect of each unique siRNA on TIIE was tested four times (data set S1), allowing the gene identities to be ranked according to their overall reduction in TIIE signal, calculated as the median of all three unique siRNA knockdowns against each gene target. The 25 cell channel genes with the greatest overall reduction in TIIE signal at 16 hpi are shown in Figure 2B. Of these, twelve were K^+^ channels; five were voltage-gated, four were from the two-pore family, two were calcium-activated and two were from the family of inward-rectifying K^+^ channels (Fig. 3). This information, taken together with the observation that four of the top five channels are involved in K^+^ conductance, suggests cellular K^+^ channels as a group are required for LCMV infection and highlights the importance of K^+^ in the LCMV multiplication cycle.

**FIGURE 3.**
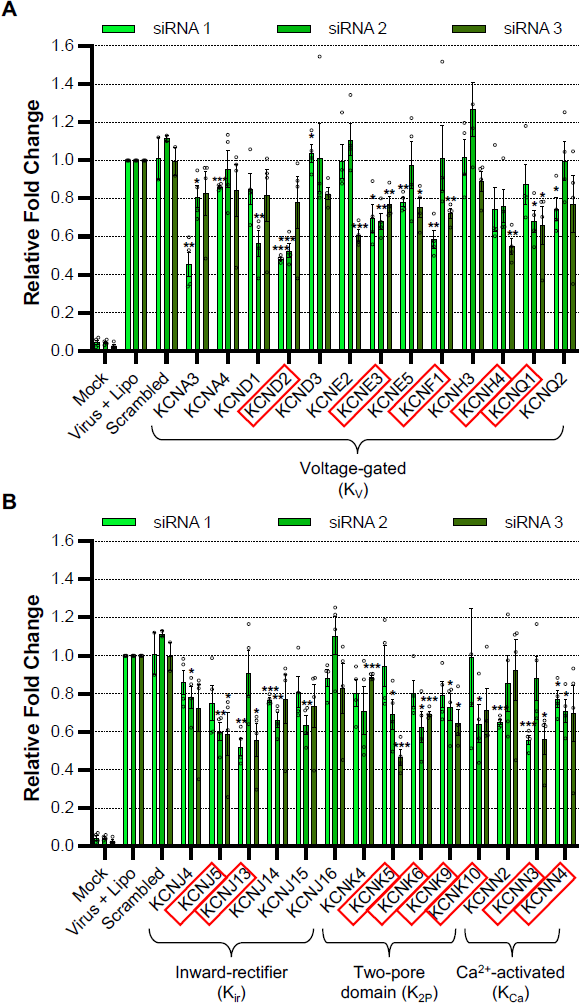
Cellular potassium ion channels are required for LCMV infection. Histograms showing the relative fold change of eGFP expression in SH-SY5Y cells after gene knockdown of a selection of potassium ion channels and infection with rLCMV-eGFP at an MOI of 0.2, after 16 hours post infection. The bars represent the knockdown eGFP expression of individual triplicate siRNAs, which targeted the same gene (siRNA 1, light green; siRNA 2, medium green; siRNA 3; dark green), as the result of four experimental repeats. Appropriate controls, such as mock-infected cells, cells infected with rLCMV-eGFP in presence of transfection reagent and cells infected with rLCMV-eGFP after treatment with scrambled siRNAs, have been included. Results were analysed by Student’s t-test whereby * = p<0.05; ** = p<0.01; *** = p<0.001. (A) shows the histograms of siRNAs targeting genes expressing voltage-gated potassium (KV) ion channels whereas (B) shows the histograms of siRNAs targeting genes expressing inward rectifier (Kir), two pore domain (K2P) and Ca2+-activated (KCa) potassium ion channels. Red boxes have been used to indicate the genes, knockdown of which, resulted in a reduction in eGFP expression below 75%.

### Pharmacological blockade of K^+^ channels prevents LCMV infection

The findings of the siRNA screen showed knockdown of several cellular K^+^ channels resulted in significantly reduced rLCMV-eGFP specific TIIE signal (Fig. 2-3). Given these data, we reasoned that blocking K^+^ channel function using pharmacological inhibitors would similarly reduce or prevent rLCMV-eGFP gene expression. To test this, we pre-treated SH-SY5Y cells with a range of concentrations of broad-range K^+^ channel inhibitors, followed by infection with rLCMV-eGFP, and measurement of TIIE signal at 24 hpi. Cell viability as an indication of compound toxicity was assessed by direct visualization of cell confluence and by MTT (3-[4,5-dimethylthiazol-2-yl]-2,5 diphenyl tetrazolium bromide) assay. To eliminate the possibility that the selected inhibitors directly influenced eGFP processing or fluorescence, the effect of channel inhibition on rLCMV-WT NP gene expression was also measured in parallel by western blot analysis, with GAPDH expression included as a loading control and to assess adverse effects of inhibitors on cell viability.

Quinidine, quinine, 4-aminopyridine (4-AP) and amiodarone all reduced rLCMV-eGFP signal and rLCMV-WT-specific gene expression levels in a dose-dependent manner at sub-toxic concentrations (Fig. 4; left and central panels). In particular, quinidine, quinine and 4-AP each reduced rLCMV-WT-specific gene expression by over 90% compared to untreated cells, at sub-toxic inhibitor concentrations, revealing high potency (Fig. 4A-C; middle panels). Amiodarone (Fig. 4D), an FDA-approved anti-arrhythmia drug with broad range K^+^ channel blocking activity, also resulted in clear reduction in rLCMV-specific gene expression at non-toxic concentrations, reducing both rLCMV-specific TIIE and NP expression by approximately 80% that of untreated infected cells at non-toxic concentrations. Inhibition of rLCMV-specific gene expression by dronedarone (Fig. 4E) and tetraethylammonium (TEA; Fig. 4F) were less convincing, showing no significant reduction in either TIIE or NP expression. Taken together, these inhibitor studies support the importance of K^+^ channels in the LCMV multiplication cycle, and further suggest that re-purposing clinically approved drugs may be a viable approach to mitigate arenavirus growth, and thus disease.

**FIGURE 4.**
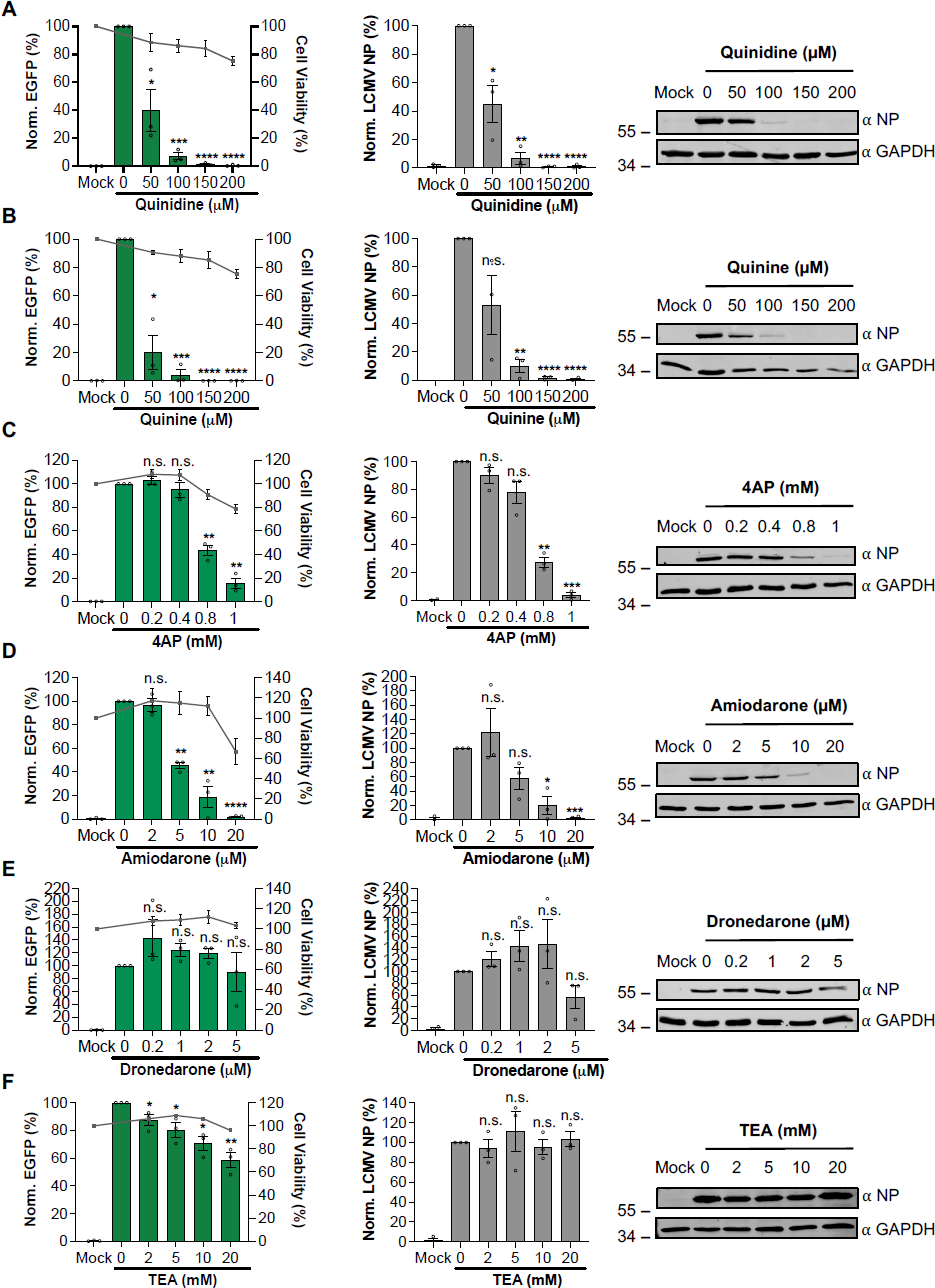
Broad-acting potassium ion channel inhibitors inhibit LCMV infection. (A-F) SH-SY5Y cells were pre-treated for 45 mins with increasing concentrations of broad-acting potassium ion channel inhibitors including tetraethylammonium (TEA; A), quinidine (B), quinine (C), 4-aminopyridine (4AP; D), amiodarone (E) and dronedarone (F). The cells were then infected in triplicate (n=3) with either rLCMV-eGFP or rLCMV-WT at a MOI of 0.1 and incubated for 24 hours. EGFP expression (left-hand panels) was measured in live cells in triplicate (n=3) using the incucyte S3 and plotted as a percentage of cells infected with rLCMV-eGFP in absence of the drug. Cell viability was also assessed using an MTS assay and plotted on the right axis. Cells infected with rLCMV-WT were lysed at 24 hours post infection and the lysates were probed for NP expression. Densitometry analysis (middle panels) was performed on the western blots in triplicate (n=3) and plotted on the graphs as a percentage of cells infected with rLCMV-WT in absence of the drug. Both eGFP expression and densitometry results were analysed by Student’s t-test whereby * = p<0.05; ** = p<0.01; *** = p<0.001, comparing results to untreated controls. Example western blots (right-hand panels) have also been included to demonstrate change in band size.

### The requirement of K^+^ channel activity acts during LCMV entry

Having established important roles for K^+^ channels in rLCMV multiplication, we next wanted to better define the stage of the infection cycle where the activity of these channels was required. To do this, we performed time-of-addition studies using the broad range K^+^ channel inhibitors quinidine, quinine and 4-AP, shown above (Fig. 4A-C) to ablate rLCMV gene expression within defined temporal windows. Cells were infected with either rLCMV-eGFP or rLCMV-WT, and then treated with each of the three inhibitors at sub-toxic concentrations at spaced time points between 0 hpi to 12 hpi, with gene expression of all infected cultures measured at 24 hpi using both western blotting with NP-antisera and live-cell eGFP detection (Fig. 5). The time points were chosen to encompass all stages of the multiplication cycle, from initial entry, to assembly and egress.

**FIGURE 5.**
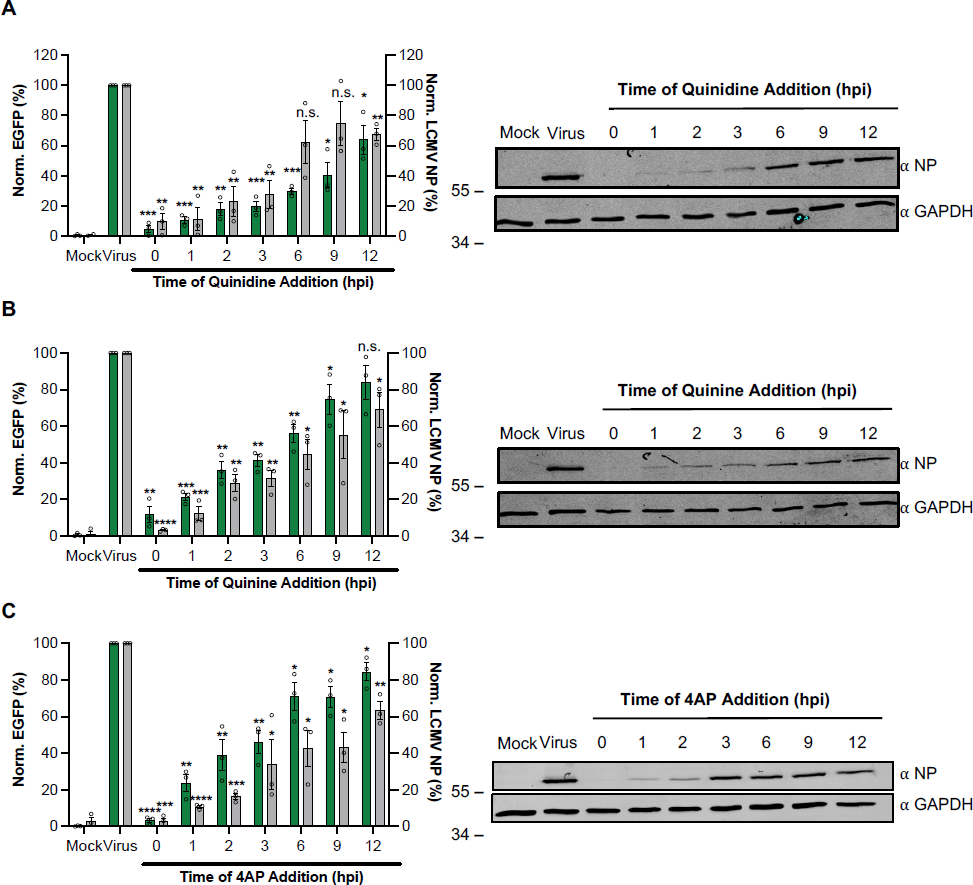
Cellular potassium ion channels are needed for early stages of LCMV infection. (A-C) SH-SY5Y cells were infected in triplicate (n=3) with either rLCMV-eGFP or rLCMV-WT at a MOI of 0.1 and incubated for 24 hours. At the indicated hours post infection (hpi; 0, 1, 2, 3, 6, 9 and 12), a suitable concentration of quinidine (A; 150 μM), quinine (B; 150 μM) or 4-aminopyridine (4AP; C; 1 mM), was added for the remainder of the incubation time. At 24 hpi, the cells were then either imaged using the incucyte S3 for analysis of eGFP expression or lysates were collected for western blot and densitometry analysis. These were plotted as a percentage of untreated EGFP or NP expression at 24 hours post infection (Virus; n=3) (left-hand panels) and example western blots have been shown (right-hand panels). Results were analysed by Student’s t-test whereby n.s. = p>0.05, * = p<0.05; ** = p<0.01; *** = p<0.001, **** = p<0.0001, comparing results to the untreated control (Virus).

In line with the pre-treatment results described above (Fig. 4A-C), quantification of western blots showed pre-treatment of quinidine at 0 hpi blocked all rLCMV-WT NP gene expression. Addition of quinidine at subsequent ‘early’ 1, 2 and 3 hpi time points reduced rLCMV-WT NP expression to near undetectable levels, whereas addition of this inhibitor at the ‘late’ time points of 6, 9 and 12 hpi had little impact. The findings were broadly similar for quinine and 4-AP, as well as corresponding TIIE measurements across all time points. Taken together, these findings suggest K^+^ channel activity is required at early stages of the LCMV replication cycle, most likely during virus internalisation and passage through the endocytic system, where we previously showed pharmacological K^+^ channel blockade collapsed K^+^ gradients^27^.

### K^+^ channel inhibition prevents LCMV endosome escape by blocking uncoating

To further examine why siRNA- and pharmacologically-mediated K^+^ channel inhibition prevented LCMV multiplication, we wanted to determine the fate of LCMV virions when infecting under conditions of K^+^ channel blockade. We postulated that K^+^ channel blockade and the subsequent collapse of endosomal K^+^ gradients prevented virions reaching a cellular destination where efficient gene expression takes place, and our goal was to determine how this occurred. To do this, we modified pUC57-LCMV-L within our rLCMV rescue system to incorporate a HA-tag into the Z matrix protein open reading frame (Supplementary Fig. 3A)^34^. This strategy provided two distinct epitopes, Z-HA and NP, within each virus particle that we could detect with specific antisera, to allow both unambiguous assignment of colocalised IF signal to infecting virions, and to allow visualisation of virus uncoating, when NP and Z proteins are proposed to spatially separate. The resulting plasmid (pUC57-LCMV-L-Z-HA) was used in subsequent rescue experiments, with generation of infectious virus bearing a HA-tagged Z protein, rLCMV-Z-HA, confirmed by detection of both NP and Z expression by western blotting in both transfected BSR-T7 and infected BHK-21 cell lysates (Supplementary Fig. 3B).

Human epithelial lung A549 cells were either untreated or pre-treated with 150 µM quinidine before being infected with rLCMV-Z-HA at an MOI of 5 and fixed at subsequent timepoints (Fig. 6). At 0 hpi, intact virions were identified on the cell surface shown by co-localisation between NP signal in red and Z-HA signal in green, similar in both untreated and quinidine-treated cells. At 3 hpi, in untreated cells, the NP signal was separated from the Z-HA signal, consistent with virion disassembly and RNP release. At 6 hpi, signals for both NP and Z were increased in intensity, with amplification of Z signal delayed relative to NP, expected due to the ambisense mode of Z gene expression. Both NP and Z adopted a distinct cellular localisation that became increasingly apparent at 9 hpi. In stark contrast, within quinidine treated cells, the NP and Z-HA signals were predominantly found co-localised in punctate regions distributed throughout the cell at all time points examined. This suggested that K^+^ channel inhibition and subsequent block of K^+^ influx had trapped incoming virions during the entry process, preventing uncoating. This further showed, for the first time, that arenavirus uncoating involves the separation of Z and NP during entry.

**FIGURE 6.**
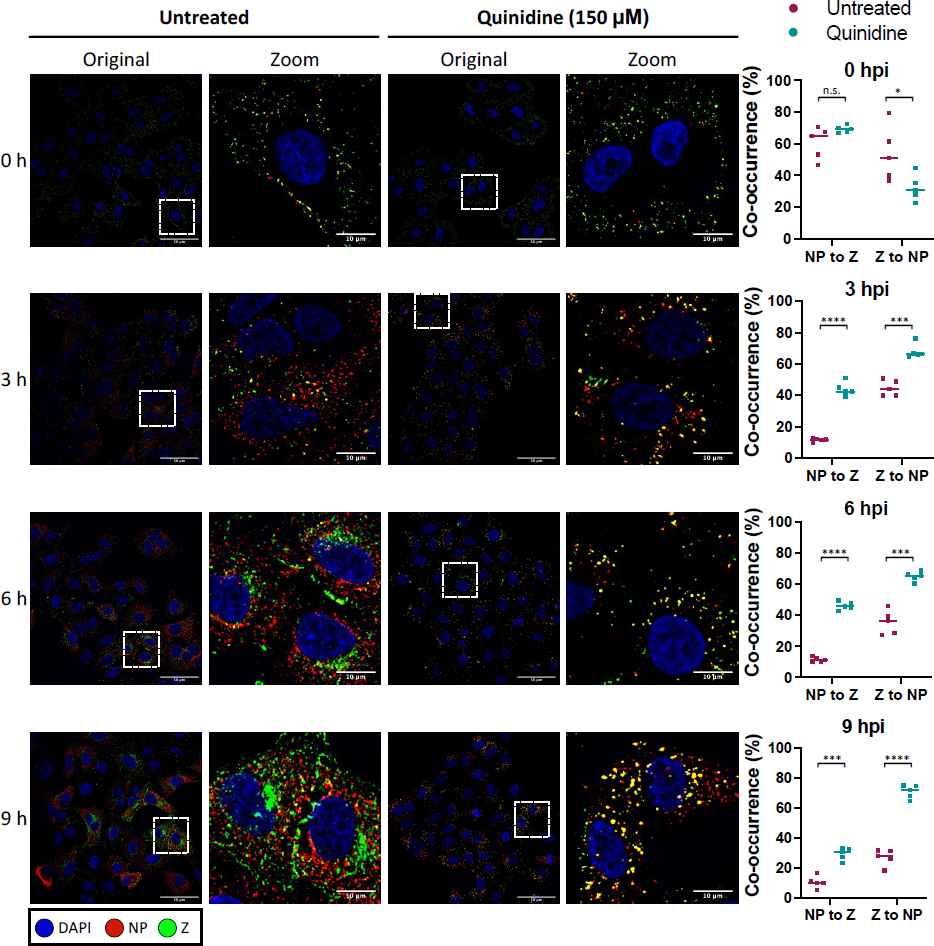
LCMV becomes trapped in presence of potassium ion channel inhibitors. A549 cells were untreated or pretreated with 150 μM quinidine and then infected with rLCMV-Z-HA at an MOI of 5 and fixed with formaldehyde at either 0, 3, 6 or 9 hours. The cells were then permeabilized, blocked and stained for the nucleus (DAPI; blue), LCMV NP (red), Z-HA (green) by indirect immunostaining. The cells were then imaged on the Olympus IX83 widefield microscope at 60 x magnification. White dashed boxes have been included to indicate which region of cells has been shown in the zoomed image. Scale bars representing 50 μm and 10 μm (zoomed images) are shown. Alongside the images is co-occurrence analysis, performed over five different images including >150 cells, which has been determined using the Manders’ coefficient method. The percentage of NP signal to Z signal and Z signal to NP signal has been compared between untreated (pink) and quinidine-treated (blue). The co-occurrence analysis between untreated and treated was further analysed through the Student’s t-test whereby n.s. = p>0.05 * = p<0.05; ** = p<0.01; *** = p<0.001; **** = p<0.0001.

### LCMV virions are trapped in late endosomes after K^+^ channel inhibition

We next sought to identify the cellular location of LCMV virions which had become trapped upon K^+^ channel inhibition. To do this, cells were either untreated or pre-treated with quinidine and infected with rLCMV-Z-HA at an MOI of 5 before fixing at 6 hpi. The 6 hpi timepoint was chosen because it represented a clear difference in the distribution of NP and Z signals between untreated and treated cells, as shown in the previous section (Fig. 6). We co-stained the cells for NP and Z, to identify intact virions, alongside cellular markers which represented different stages of the endocytic pathway: Rab 5 for early endosomes, Rab 7 for late endosomes, LAMP1 for lysosomes and CD164 as an established LCMV late endosomal secondary receptor. Co-occurrence analysis was then calculated on over 150 cells across five images. Then, to examine only points at which both the NP and Z signals were present, indicative of specific virions, the NP and Z signals were multiplied together (NP/Z), and the Manders’ overlap coefficient was calculated to determine what percentage of the NP/Z signal overlapped with the cell marker signal.

Our results showed that when cells were treated with quinidine to block endosomal K^+^ influx, there was a significant increase in the co-occurrence between NP/Z and Rab7 signals, resulting in approximately 70 % of signal overlap (Fig. 7; second row of panels). A similar trend was also found when the co-occurrence analysis was performed on CD164 signal (∼ 70 % overlap between NP/Z and CD164 signals; Fig. 7; fourth row of panels). This suggested that there was a significant accumulation of trapped virions in Rab7-positive and CD164-positive late endosomes after quinidine treatment. Little co-occurrence was seen between NP/Z signal and that of Rab5 (∼ 20 %) and there was no significant increase in co-occurrence after quinidine treatment (Fig. 7; first row of panels). Furthermore, less than 5 % of the NP/Z signal co-occurred with LAMP1 signal after quinidine treatment (Fig. 7; third row of panels). Taken together, these data suggest that when endosomal K^+^ influx is blocked through K^+^ channel inhibition, virions progress through Rab5-positive early endosomes, become trapped in Rab7/CD164-positive late endosomes and do not reach LAMP1-positive lysosomes. Thus, LCMV RNPs cannot escape from the trapped virion, gene expression cannot occur, and infection is blocked.

**FIGURE 7.**
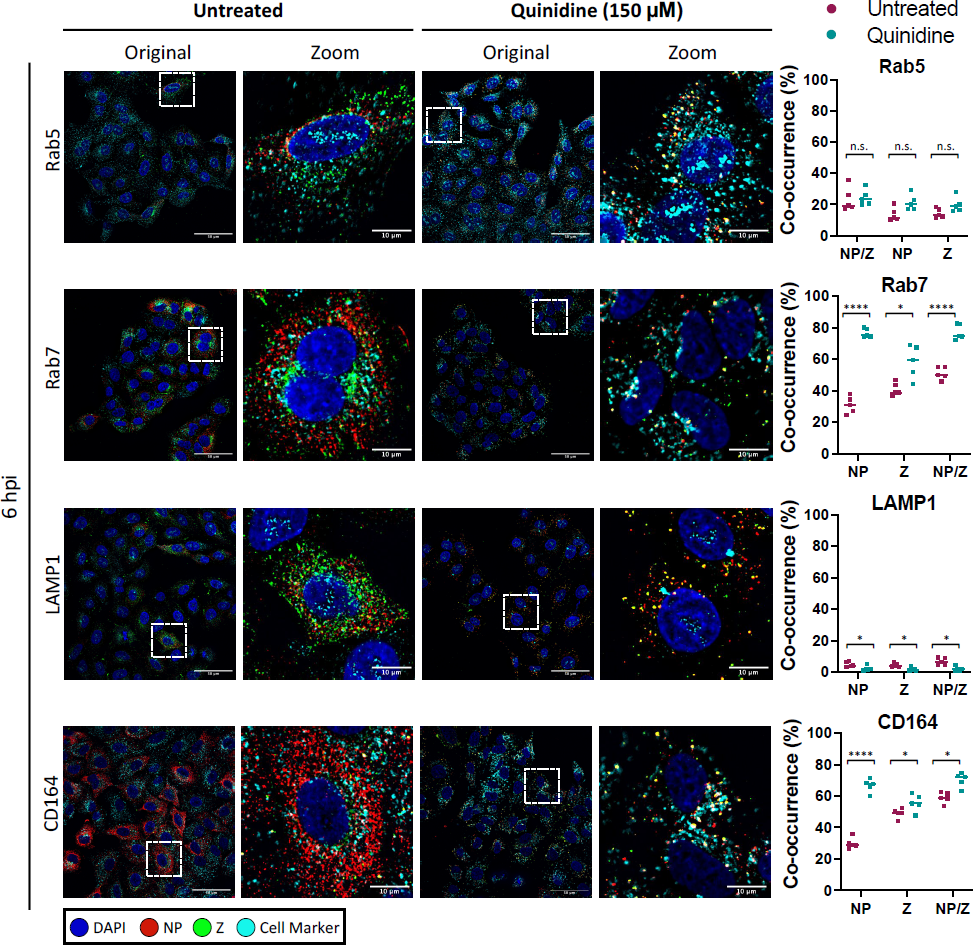
LCMV NP puncta colocalise with Rab7 and CD164 in cells treated with potassium ion channel inhibitors. A549 cells were untreated or pretreated with 150 μM quinidine and then infected with rLCMV-Z-HA at an MOI of 5 and fixed with formaldehyde at 6 hours post infection (hpi). The cells were then permeabilised, blocked and stained for the nucleus (DAPI; blue), LCMV NP (red), Z-HA (green) and for cellular markers (cyan), Rab5, Rab7, CD164 and LAMP1, by indirect immunostaining. The cells were then imaged on the Olympus IX83 widefield microscope at 60 x magnification. White dashed boxes have been included to indicate which region of cells has been shown in the zoomed image. Scale bars representing 50 μm and 10 μm have been included. Alongside the images is co-occurrence analysis, performed over five different images including >150 cells, which has been determined using the Manders’ coefficient method. The percentage of either NP signal, Z signal or NP and Z signal has been examined against the signal of the marker and comparison between untreated (pink) and quinidine-treated (blue) was further analysed through the Student’s t-test whereby n.s. = p>0.05, * = p<0.05; ** = p<0.01; *** = p<0.001; **** = p<0.0001.

### Exposure to high K^+^ ion concentration reduces GP1 binding to CD164

The results of the previous section showed K^+^ channel blockade led to entrapment of LCMV in late endosomes by preventing uncoating, suggesting K^+^ is required for virion uncoating, and we are curious as to why this might be.

Late endosomal compartments supply the two known requirements for LCMV fusion and escape, namely low pH (5.4) and abundant CD164. Despite this, under K^+^ channel blockade, LCMV becomes trapped and unable to uncoat. As the GP1/CD164 interaction at low pH alone (with no additional K^+^) is known to promote membrane fusion^19^, we reasoned K^+^ may exert its influence on uncoating by one of two options; either by modulating the interaction between external virion component GP1 and its CD164 ligand, or by influencing separation between internal virion components such as between the RNP and Z (i.e. uncoating). This latter possibility is particularly intriguing considering a similar mechanism proposed for influenza virus^26^ and the recent demonstration that the arenavirus interior is accessible to ion influx via formation of a glycoprotein-associated envelope pore in response to low pH^35, 36^.

To test whether K^+^ modulated the GP1/CD164 interaction, we utilized a previously described ELISA-based binding assay^19^, where immobilized LCMV GP1 was incubated with soluble CD164 at endo-lysosomal relevant pH values between 5.8 and 4.8, either in the presence or absence of K^+^. Interestingly, we found GP1/CD164 binding curves were similar irrespective of K^+^ concentration, with maximal binding in both cases occurring at pH 5.4 (Fig. 8) consistent with the late-endosomal site of LCMV entry. This suggests the critical role of K^+^ in uncoating is not to promote the GP1/CD164 interaction. Based on our results, and those of others, we suggest K^+^ influx to the virion interior influences interactions between internal Z matrix and RNPs (Fig. 9)^26, 35, 36^.

**FIGURE 8.**
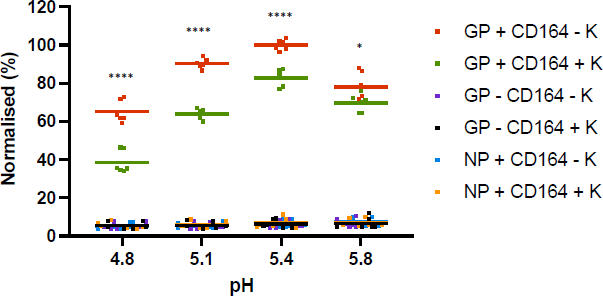
LCMV GP1 binding to CD164 is reduced in presence of potassium. Soluble LCMV GP1 was used to coat 96-well high-protein binding plates overnight and then the wells were washed and blocked before washing in buffers at the specified pH, containing either 5 mM potassium chloride (GP + CD164 – K; red) or 140 mM potassium chloride (GP + CD164 + K; green). CD164-Fc was then added at the specified pH, washed and incubated with goat anti-human IgG-HRP. The wells were washed and TMB substrate was added and the reaction was stopped with sulphuric acid, before reading the plate at 450 nm on a spectrophotometer plate-reader. The experiment was performed three times in duplicate and all results have been shown. All results were normalised to GP + CD164 – K at pH 5.8, which was condition that resulted in the strongest absorption. Controls whereby LCMV NP was bound to the plate instead of GP1 (NP + CD164 – K; blue or NP + CD164 + K; orange) or where no CD164 was added (GP – CD164 – K; purple or GP – CD164 – K; black) were included to show lack of binding between NP and CD164 or between GP1 and goat anti-human IgG-HRP. The difference between 5 mM potassium chloride or 140 mM potassium chloride at the respective pH was further analysed through the Student’s t-test whereby n.s. = not significant; * = p<0.05; ** = p<0.01; *** = p<0.001; **** = p<0.0001.

**FIGURE 9.**
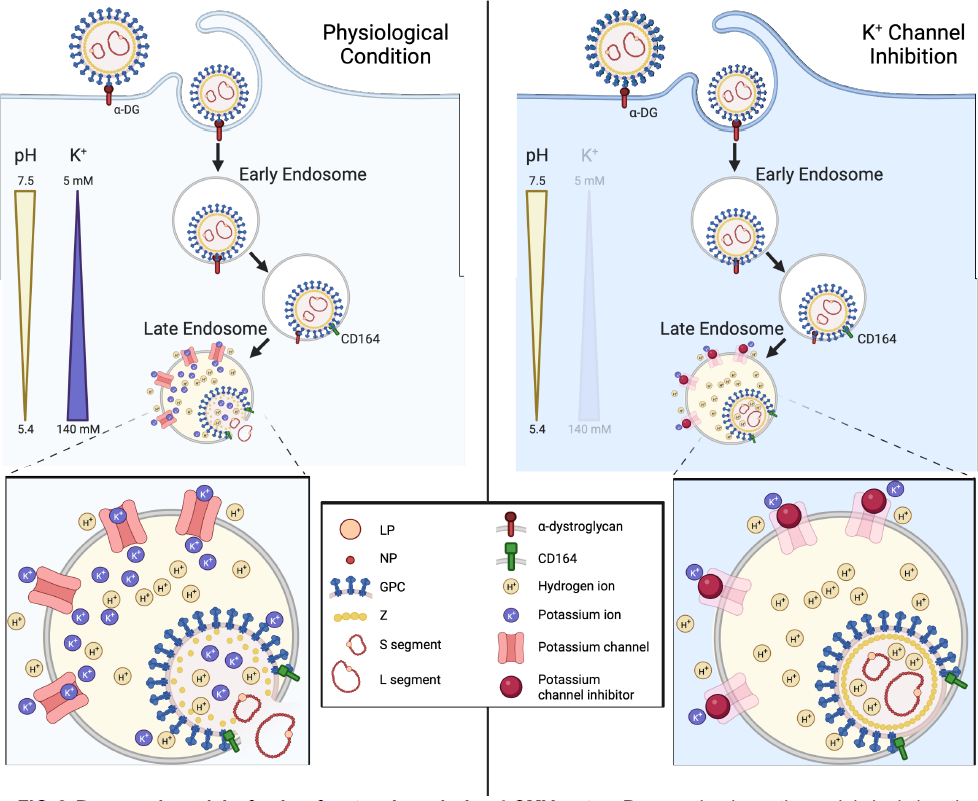
Proposed model of the role of potassium during LCMV entry. Proposed schematic model depicting the entry pathway of LCMV under physiological conditions and under conditions of potassium (K+) depletion. LCMV is internalised through macropinocytosis, after binding to alpha-dystroglycan (ɑ-DG), into an early endosome. LCMV then progresses through the endocytic system to late endosomes where it experiences a reduction in pH and an increase in K+ concentration. The low pH is sufficient to drive a receptor switch to secondary receptor CD164, and the low pH and CD164 interaction drives membrane fusion and RNP release. We propose that in addition to hydrogen ions, K+ ions are transported into the virion interior to mediate destabilisation and subsequent uncoating of the Z matrix layer. Under conditions of K+ depletion, we see incoming virions trapped in Rab7+ and CD164+ late endosomes, but we suggest that the absence of K+ in the endosome means no K+ is present in the virion interior, preventing uncoating of the viral matrix layer and release of the RNPs.

## DISCUSSION

During viral entry to host cells, viruses encounter a complex environment with drastic changes in ionic concentration and membrane composition along the endocytic pathway^37^. This includes a reduction in pH, concurrent with an increase in the concentrations of other ions. Previously, our group has demonstrated a role for K^+^ ions in the entry process of HAZV^28^, when K^+^ induces structural changes in the external viral glycoprotein spikes to form a fusion intermediate conformation. Additionally, for BUNV, we showed K^+^ has a direct effect on virion morphology GP architecture and function, beyond that elicited by acidic pH, with effects confined to the floor region of the GPC complex^30^. The requirement of K^+^ for orthobunyavirus entry has further been demonstrated by others for La Crosse virus, Keystone virus and Germiston virus^38, 39^.To extend this work across the *Bunyavirales* order, we examined the role of K^+^ flux in the multiplication cycle of LCMV, the prototypic member of the *Arenaviridae* family. Using an eGFP-expressing LCMV variant generated through manipulation of an infectious clone, alongside both a siRNA screen targeting human ion channels and K^+^ ion channel inhibitors, we showed LCMV infection depended on K^+^ ion channel function. Using recombinant LCMV bearing epitope-tagged components, we further pinpointed the role of K^+^ to the LCMV entry process, with K^+^ channel blockade resulting in entrapment of intact virions within late endosomes. Based on these results, we propose a model whereby K^+^ is necessary for virion uncoating, which represents the process when internal RNP are released from the virion interior. Specifically, we propose this involves dissociation between RNPs and the Z matrix layer, in a similar way as has been described for influenza virus^26^. Our demonstration that LCMV NP and Z separate during entry, whereas remain associated under K^+^ channel blockade, suggest that RNP release into the cytosol involves dissociation from the Z matrix layer, and to the best of our knowledge, this represents the first time this has been shown.

Our model (Fig 9) seeks to explain our findings, alongside those of others. In our current study, we showed that K^+^ channel blockade, thus preventing endosomal K^+^ influx, arrested LCMV in late endosomes where its two known entry requirements, namely CD164 and fusion-enabling low pH of around 5.4, are encountered. Consequently, the late endosome block in LCMV entry under K^+^ depleted conditions is perplexing but suggests K^+^ represents a third requirement for LCMV endosome escape, in addition to low pH and CD164.

In broad terms, endosome escape involves at least two distinct stages; a GPC-mediated fusion step, and an uncoating step, where RNPs exit the confines of the virion and are released into the cytosol, although whether these steps are sequential or concurrent is unknown. Interestingly, Bakkers *et al* (2022) used a pseudovirus fusion assay to show that LCMV GP1 binding CD164 at low pH is sufficient for the first envelope fusion step^19^. This, combined with our findings, points towards a role for K^+^ in the virion uncoating step, which we depict in our model (Fig. 9).

K^+^ mediated virion uncoating is established for influenza virus, which involves K^+^ influx into the virion interior, facilitated by the M2 transmembrane viroporin^26^. For influenza virus, K^+^ influx results in the dissociation of the matrix layer M1 that encases the eight RNP segments, which are then released into the cytosol for subsequent trafficking to the nucleus. In contrast, arenaviruses do not express a dedicated viroporin component as a functional analog of M2, and so the question of how K^+^ might penetrate the LCMV envelope to influence interior components is highly pertinent.

Interestingly, recent work suggests that the arenavirus envelope can be breached by H^+^ ions, resulting in acidification of the virion interior, measured using pH sensitive indicator dyes in the context of lentiviral based pseudovirus particles^35^. Further to this, it has been proposed that arenavirus envelope permeability is mediated by structural rearrangements of the arenavirus glycoproteins themselves, following interactions with cellular factors at low pH^36^. These structural rearrangements are proposed to lead to the formation of a transmembrane pore, allowing the passage of H^+^ into the virion interior as a critical entry trigger promoting virion uncoating. This model is similar to ours, with the additional requirement that K^+^ virion influx is needed for the uncoating step. Currently, the question of whether such a glycoprotein pore can allow the influx of K^+^ is untested, although as for influenza virus M2, it is reasonable to suggest that movement of ions might not be so selective. The functional parallels that necessarily exist between the entry of influenza virus and arenaviruses are interesting, due to their shared possession of a matrix layer, which must be disassembled before RNPs can be released; and the fact that both viruses share viral fusion proteins of similar overall architecture^40^. The results and model presented here suggest that the solution to this internal disassembly barrier has been solved in a similar way, using viroporin-like activity.

This work supports a critical role for K^+^ ions in the LCMV replication cycle and represents the first study to establish the identity of cellular K^+^ channels that play a role in the arenavirus replication cycle. Interestingly, a previous report identified TRAM34 as a potent inhibitor of arenavirus entry, pointing towards a role for the TRAM34 therapeutic target, namely the cellular calcium-activated K^+^ channel KCNN4 (KCa3.1)^41^. However, the anti-arenaviral activity of TRAM34 was found to be independent of the ion channel target, and while the compound was found to act at the fusion stage of virus entry, the underlying mechanism of its activity remains unknown. It will be interesting to determine which of the several K^+^ channels that were identified in the LCMV siRNA screen are key contributors to endosomal K^+^ influx, with the aim of understanding how K+ influences both cellular and viral processes, as well as identifying host cell targets for antiviral compounds. Since, cellular K^+^ channels are tested targets for clinically approved human therapies, this raises the possibility that current licensed FDA-approved K^+^ ion channel inhibitors could be repurposed for effective management of LCMV infection, or by extrapolation, to treat more serious arenavirus infections such as Lassa fever caused by LASV.

## MATERIALS AND METHODS

### Cell Lines and Virus

BHK-21 cells, BSR-T7 cells, SH-SY5Y cells and A549 cells were acquired from ATCC and maintained in high-glucose Dulbecco’s modified Eagle medium (DMEM; Sigma-Aldrich), supplemented with 10 % heat-inactivated foetal bovine serum (FBS), 100 µg of streptomycin/mL and 100 U of penicillin/mL, and incubated in a humidified incubator at 37 °C with 5 % CO_2_. BSR-T7 cells constitutively expressing T7 RNA polymerase (T7P)^42^ were also additionally supplemented with G418 (500 µg/mL) every other passage to maintain the T7P plasmid. LCMV strain Armstrong virus and sequences were obtained from the National Collection of Pathogenic Viruses (NCPV; 1805291v), managed by the UK Health Security Agency.

### Plasmids

Full-length cDNAs representing the intact S and L segments were designed as previously reported^43, 44^ and synthesised (GENEWIZ) using the LCMV strain Armstrong: clone 13 derivative (GenBank accession numbers: DQ361065 and DQ361066 for S and L respectively) in pUC57, subsequently named pUC57-S and pUC57-L.

Additional supporting plasmids pUC57-NP and pUC57-LP were constructed using PCR amplification to sub-clone the NP ORF and the LP ORF from pUC57-S and pUC57-L, respectively, into the pUC57 vector using complementary flanking restriction sites.

Plasmid pUC57-S-eGFP was synthesised by sub-cloning the eGFP-P2A ORF (GenBank accession number: QFU20120.1; GENEWIZ) into pUC57-S, using complementary flanking restriction sites. The eGFP-P2A ORF was inserted between the start codon of the NP ORF and the 3’ end of the untranslated region (UTR) of the S segment. The eGFP ORF was separated from the NP ORF by the porcine teschovirus 1 (PTV-1) 2A self-cleaving peptide sequence (ATNFSLLKQAGDVEENPGP) (GenBank accession number: MH358390). Plasmid pUC57-L-Z-HA was synthesised through PCR insertion of the HA tag (YPYDVPDYA) at the end of the Z ORF, prior to the terminal stop codon. All primers sequences are available upon request.

### Generation of recombinant LCMV from cDNA

BSR-T7 cells were seeded in 6-well plates at a density of 2×10^5^ cells/well. Following overnight incubation, the cells were transfected with 1.6 µg of pUC57-S or pUC57-S-eGFP, 1.6 µg of pUC57-NP, 2.8 µg pUC57-L or pUC57-L-Z-HA, 2 µg of pUC57-LP and 0.6 µg of pUC57-T7 in 200 µL OptiMEM media, followed by 2.5 µl/µg TransIT-LT1 transfection reagent (Mirus). Control transfections were also performed, in which pUC57-L and pUC57-LP were excluded. At 24-hour post transfection (hpt), the media was removed and replaced with 1 mL of DMEM supplemented with 2 % FBS. At 120 hpt, the supernatants were collected, clarified and used to infect BHK-21 cells, seeded the previous day at 2×10^5^ cells/well in 6-well plates.

### Focus forming assay

Focus forming assays were performed on confluent monolayers of BHK-21 cells. Serial dilutions of LCMV (10-fold) made in serum-free DMEM were added to the cells and incubated for 1 h at 37 °C. The dilutions were removed from the cells and overlaid with 0.8 % methylcellulose. After 3 days incubation at 37 °C, the overlay was removed and the cells were fixed with 4 % formaldehyde in PBS, permeabilised with 0.3 % Triton-X-100 and blocked using 1 % bovine serum albumin (BSA). The cells were then incubated for 1 h with anti-LCMV NP polyclonal antibody (1:1000), made in 1 % BSA. The cells were then stained for 1 hour at room temperature with an anti-sheep Alexa Fluor secondary antibody (Cell Signaling Technologies). The entire well was then imaged at 4X magnification using the IncuCyte S3 automated microscope (Sartorius). Foci were counted and the titre was determined.

### Western Blot Analysis

Cells were lysed with 1X radioimmunoprecipitation assay (RIPA) buffer (50 mM Tris-HCl pH 7.5, 150 mM NaCl, 1 % (v/v) NP40 alternative, 0.5 % (w/v) sodium deoxycholate and 0.1 % sodium dodecyl sulphate (SDS; w/v)), supplemented with 1X cOmpleteTM, Mini, EDTA-free Protease I inhibitor cocktail (Sigma-Aldrich) for 15 minutes on ice. Lysates were collected, proteins were resolved on a 12 % SDS-PAGE gel and then transferred to a polyvinylidene fluoride (PVDF) membrane. The transfer was performed at 15 V for 30 min, using the Trans-Blot turbo (Bio-Rad). After transfer, the membrane was blocked for 1 hour in Odyssey blocking buffer (PBS) (Licor; diluted 1 : 1 with 1X PBS). Subsequently, the membrane was stained with the primary antibodies (made in 1:4 [blocking buffer:PBS-T]; anti-LCMV NP 1:1000, anti-GAPDH 1:7500, anti-GFP (B-2) 1:1000, anti-HA Tag 1:1000) for 1 hour rocking, at room temperature and then with corresponding secondary antibodies (made in 1:4 [blocking buffer:PBS-T]; all secondary antibodies used at 1:10000) for 1 hour at room temperature. The membrane was dried and visualised on the Licor Odyssey Sa Infrared imaging system. Densitometry analysis was performed using ImageJ over three independent experiments.

### Assessment of Viral Release

SH-SY5Y cells (5×10^4^ cells/well) were seeded into a 48-well plate and incubated overnight. The cells were then washed with PBS and infected with rLCMV-eGFP at an MOI of 0.2 for 1 hour at 37 °C. After the incubation period, the infecting media was removed and washed four times with PBS, before adding DMEM (supplemented with 2.5% FBS). At given timepoints, supernatant from infected cells was removed and added to fresh cells (transfer cells). This was also performed for cells that were mock-infected. The original infected cells were imaged for eGFP expression with the incucyte S3 to confirm all wells had been infected with an equal amount of virus. The transfer wells were imaged for eGFP expression with the incucyte S3 every hour for 24 hours following the last time point. Analysis was performed on the images taken at 24 hours after the supernatant from the infected cells was added to the transfer cells.

### RNAi Screen

A master mix of 0.3 µL of Lipofectamine RNAiMAX reagent (Invitrogen) and 16.7 µL of Opti-MEM per well was pipetted into each well of a 96-well plate. Each siRNA from the 33 Silencer™ Human Ion Channel siRNA Library (Invitrogen) was transferred into the Opti-MEM mixture, at a final concentration of 3 pmol. This was incubated for 20 min, after which 100 µL of a cell suspension of 2 × 10^5^ SH-SY5Y cells/mL in 10 % DMEM was added to each well. Cells were incubated with the transfection and siRNA mix for 24 h at 37 °C, following which 60 µL of the medium was removed and replaced with 200 µL of fresh 10% FBS DMEM to dilute the siRNAs and transfection reagent. After a further 6 h, the medium was removed, and cells were washed in PBS, infected with rLCMV-eGFP at an MOI of 0.2 and incubated at 37 °C until 16 hpi when the eGFP expression was analysed using the Incucyte S3. The total integrated intensity of eGFP (TIIE; green count units [GCU] × μm*2*/image) was first normalised to confluency per well and then analysed as a percentage of the total green integrated intensity in positive control wells containing virus and lipofectamine only. Normalised values were averaged between two technical repeats and two biological repeats.

### Infection Drug Assays

1 × 10^5^ SH-SY5Y cells/well were seeded into 12-well plates and incubated overnight. Cells were pre-incubated for 45 mins with media supplemented with the specific ion channel inhibitor or the solvent control of each inhibitor. Cells were then incubated for 1 hour with rLCMV-eGFP or rLCMV-WT at an MOI of 0.1, which was then replaced with the media from the pre-incubation. The cells were then incubated for a further 24 hours before analysis of eGFP expression or NP expression by Western blot analysis.

### Timecourse of Drug Addition

LCMV-infected SH-SY5Y cells were treated with quinidine (150 μM), quinine (150 μM) or 4AP (1 mM) during the 24 h infection period (T = 0), or added 1, 2, 3, 6, 9, and 12 hpi. Infection was allowed to proceed for a total of 24 h, and cells were analysed for eGFP expression or NP expression by Western blot analysis.

### Immunofluorescence Analysis - Widefield

To assess localisation of LCMV proteins during entry, rLCMV-Z-HA-infected A549 cells were grown on glass coverslips. At specified timepoints, the cells were fixed with 4 % formaldehyde for 10 mins, permeabilised with 0.3 % triton-x-100 for 10 mins and blocked with 1% bovine serum albumin (BSA) for 1 hour. Cells were then labelled with anti-LCMV NP (1:500), anti-HA (for Z staining; 1:500) and antibodies against cell markers (Rab5; 1:200, Rab7; 1:100, LAMP1; 1:100, CD164; 1:200) for 1 hour, followed by labelling with corresponding Alexa Fluor 594 nm, 488 nm or 647 nm secondary antibodies (all secondary antibodies used at 1:500), respectively, for 1 hour. Cells were then washed and mounted onto microscope slides using prolong gold containing DAPI (Invitrogen-Molecular Probes). Stained cells were viewed on the Olympus IX83 widefield microscope at 60 x magnification. Z stack images were collected and the slice showing the strongest signal was chosen for the display image. Five randomly chosen fields were imaged for each condition, with a minimum of 20 cells in each field, to provide statistical co-occurrence analysis of >100 cells. Co-occurrence analysis was performed using an Otsu threshold generated in ImageJ software and the Manders’ coefficient method was performed to determine co-occurrence.

### LCMV pH/ion assays

LCMV (3 μL; MOI = 0.01) was mixed with high potassium buffers (30 µL; 140 mM potassium chloride) at pH 7.3, 6.3, 5.4, 5.2 or 5.0 at a 1:10 ratio and was incubated at 37 °C for either 2 hours or 5 mins. Control experiments were carried out by incubation in buffers at pH 7.3, 6.3, 5.4, 5.2 or 5.0 with low potassium concentration (5 mM potassium chloride). The priming buffers were then diluted out at a ratio of 1:50 and immediately added to SH-SY5Y cells. The cells were incubated for a further 18 hours before lysis and Western blot analysis.

All buffers contained 12 mM sodium chloride and either 5 mM (low) or 140 mM (high) potassium chloride. The buffering agent for the buffers changed depending on pH; for pH 7.3 20 mM tris was used, for pH 6.3 30 mM bis-tris was used and for pHs 5.4-5.0 50 mM sodium citrate was used. The pHs of the buffers were adjusted on the day of the experiment, using hydrochloric acid for pH 7.3 and 6.3 buffers, and citric acid for pH 5.4-5.0 buffers.

### Enzyme-Linked Immunosorbent Assay

The enzyme-linked immunosorbent assay (ELISA) used here was adapted from Bakkers et al., 2022. High protein-binding MaxiSorp plates (ThermoFisher Scientific) were coated with 200 ng/well soluble LCMV sGP resuspended in 1x TBS and incubated overnight at room temperature. Wells were then washed with 1x TBS and incubated with 3% BSA at 37 °C for 1 hour. Wells were extensively washed in ELISA buffer composed of 0.1 M citrate buffer, 3% BSA, 0.05% Tween 20, 1 mM calcium chloride, 1 mM magnesium chloride, 150 mM sodium chloride and either 5 mM or 140 mM potassium chloride at the respective pHs. 200 ng/well soluble Fc-tagged CD164 in respective ELISA buffers was added and incubated at 37 °C for 1 hour. Wells were then extensively washed in ELISA buffer and incubated with goat anti-human IgG conjugated to horseradish peroxidase (HRP; 1:500) in the respective ELISA buffers for 1 hour at 37 °C. Wells were once again washed extensively with ELISA buffer at the respective pH. TMB (3,3′,5,5′-tetramethylbenzidine) ELISA Substrate (ThermoFisher Scientific) was added for 15 mins, followed by equal volumes of 2 M sulphuric acid to stop the reaction. The optical density at 450nm (OD450) of each well was read using a spectrophotometer plate reader.

### Statistical Analysis

The statistical significance of data were determined by performing a Student’s *t* test. Significance was deemed when the values were less than or equal to the 0.05 *p* value.

## Supporting information

supplemental siRNA dataset

## ACKNOWLEDGEMENTS

We acknowledge funding from MRC grant MR/T016159/1 in support of AS and HNT, BBSRC PhD studentship grant to AS, UKHSA PhD studentship grant to OB. The authors thank Dr Ruth Hughes and Dr Sally Boxall of the bioimaging facility, faculty of biological sciences, University of Leeds for their expert assistance and use of the Zeiss LSM880, funded by Wellcome Trust grant WT104918MA

---

**SUPPLEMENTAL FIGURE S1.**
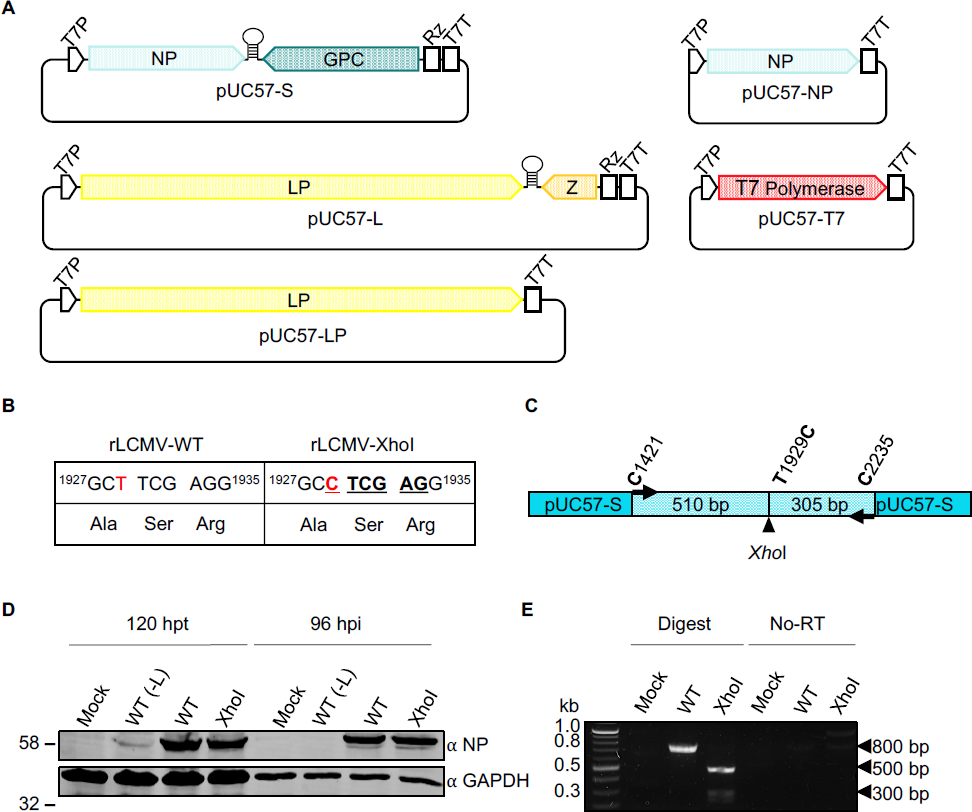
Development of reverse genetics system to generate recombinant LCMV. (A) Schematic depiction of the plasmids encoding the S segment and L segment of LCMV, in addition to the support plasmids encoding the nucleocapsid protein (NP), the RNA-dependent RNA-polymerase (LP) and the T7 polymerase (T7). Also shown flanking the LCMV S and L segments are the T7 polymerase promoter (T7P), hepatitis delta virus ribozyme (Rz) and the T7 polymerase terminator (T7T) sequences. (B) A silent mutation was introduced to show that the rLCMV-WT derived from the reverse genetics plasmids. T1929 (red) was mutated to C (red) introducing a XhoI restriction site (underlined) without disrupting the amino acid sequence. (C) Primers were designed to bind 510 base pairs upstream (C1421) and 305 bp downstream (C2235) from the T1929C mutation in the pUC57-S plasmid. (D) Lysates were collected from BSR-T7 cells transfected with pUC57-L, pUC57-LP, pUC57-NP, pUC57-T7 and either pUC57-S (WT) or pUC57-S-XhoI (XhoI) at 120 hours post transfection and from BHK-21 cells infected with the supernatant from the BSR-T7 cells, 96 hours post infection. Appropriate controls were included whereby the L-expressing plasmids were omitted to prevent rescue of infectious virus (-L). The lysates were probed for NP expression, and GAPDH as a loading control. (E) Supernatant from the infected BHK-21 cells was used to infect fresh BHK-21 cells and at 96 hpi, the supernatant was harvested and RNA was extracted. The extracted RNA was reverse transcribed into complementary DNA and digested with XhoI before agarose gel electrophoresis analysis. A control experiment lacking the reverse transcription step was performed, demonstrating the bands were not derived from any contaminating DNA plasmids carried over from the transfected cells.

**SUPPLEMENTAL FIGURE S2.**
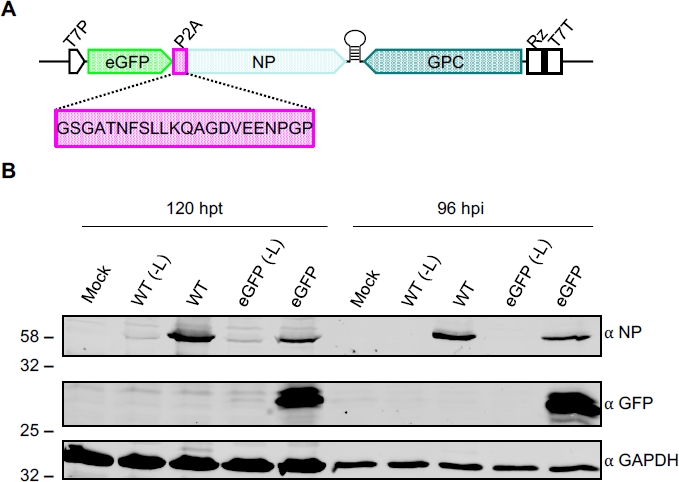
Generation of an eGFP-expressing variant of LCMV. (A) Schematic depiction of plasmid pUC57-LCMV-S encoding the S segment of LCMV, with the insertion of the eGFP open reading frame upstream of the NP gene. The eGFP and NP genes are separated by a porcine teschovirus 2A linker (P2A), the amino acids of which has been shown in the pink box. Also depicted flanking the LCMV S segment are the T7 polymerase promoter (T7P), hepatitis delta virus ribozyme (Rz) and the T7 polymerase terminator (T7T). (B) Lysates were collected from BSR-T7 cells transfected with the LCMV rescue plasmids 120 hours post transfection (hpt), and from BHK-21 cells infected with the supernatant from the BSR-T7 cells at 96 hours post infection (hpi). Appropriate controls were included whereby the L-expressing plasmids were omitted to prevent rescue of infectious virus (-L). The lysates were probed for NP and EGFP expression, and GAPDH as a loading control using specific antisera, as indicated.

**SUPPLEMENTAL FIGURE S3.**
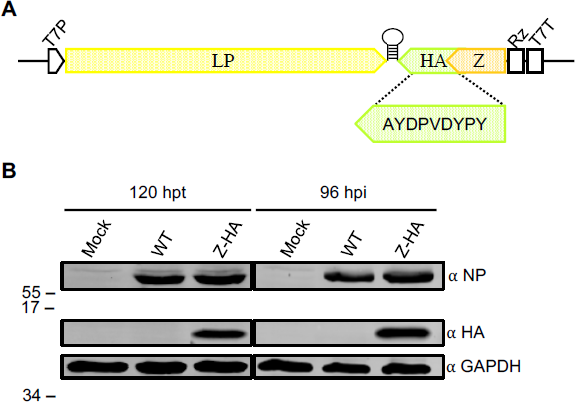
Rescue of a variant of LCMV expressing a HA-tagged Z. (A) Schematic depiction of plasmid pUC57-LCMV-L-Z-HA encoding the L segment of LCMV, with the insertion of the HA tag (the amino acids of which has been shown in the green box) downstream of the Z open reading frame. Also depicted flanking the LCMV L segment are the T7 polymerase promoter (T7P), hepatitis delta virus ribozyme (Rz) and the T7 polymerase terminator (T7T). (B) Lysates were collected from BSR-T7 cells transfected with the reverse genetics plasmids 120 hours post transfection (hpt) and from BHK-21 cells infected with the supernatant from the BSR-T7 cells, 96 hours post infection (hpi). rLCMV-WT was recovered alongside rLCMV-Z-HA. The lysates were probed for NP and HA expression, and GAPDH as a loading control.

